# The Cf-4 receptor-like protein associates with the BAK1 receptor-like kinase to initiate receptor endocytosis and plant immunity

**DOI:** 10.1101/019471

**Authors:** Jelle Postma, Thomas W. H. Liebrand, Guozhi Bi, Alexandre Evrard, Ruby R. Bye, Malick Mbengue, Matthieu H. A. J. Joosten, Silke Robatzek

## Abstract

- The first layer of plant immunity is activated by cell surface receptor-like kinases (RLKs) and proteins (RLPs) that detect infectious pathogens. Constitutive interaction with the RLK SUPPRESSOR OF BIR1 (SOBIR1) contributes to RLP stability and kinase activity. As RLK activation requires transphosphorylation with a second associated RLK, it remains elusive how RLPs initiate downstream signaling. To address this, we investigated functioning of Cf RLPs that mediate immunity of tomato against *Cladosporium fulvum*.
- We employed live-cell imaging and co-immunoprecipitation in tomato and *Nicotiana benthamiana* to investigate the requirement of associated kinases for Cf activity and ligand-induced subcellular trafficking of Cf-4.
- Upon elicitation with the matching effector ligands Avr4 and Avr9, BRI1-ASSOCIATED KINASE 1 (BAK1) associates with Cf-4 and Cf-9. Furthermore, Cf-4 that interacts with SOBIR1 at the plasma membrane, is recruited to late endosomes after elicitation. Significantly, BAK1 is required for Avr4-triggered endocytosis, effector-triggered defenses in Cf-4 plants and resistance of tomato against *C. fulvum*.
- Our observations indicate that RLP-mediated immune signaling and endocytosis require ligand-induced recruitment of BAK1, reminiscent of BAK1 interaction and subcellular fate of the FLAGELLIN SENSING 2 RLK. This reveals that diverse classes of cell surface immune receptors share common requirements for signaling initiation and endocytosis.

## Introduction

The innate immune system of higher eukaryotes senses a wide range of pathogens through distinct pattern recognition receptors (PRRs). The engagement of these receptors leads to the activation of signaling pathways resulting in the induction of defense responses. Plants rely on plasma membrane-resident receptor-like kinases (RLKs) and receptor-like proteins (RLPs) as a first layer of the immune system against phytopathogenic microbes, many of them exploiting the extracellular space of plant tissues for their growth (Faulkner & Robatzek, 2012). Therefore, decoding the mechanisms underlying the perception of pathogen-derived signals by RLKs and RLPs at the host cell surface and subsequent activation of defense signaling is an important aspect for understanding how plant immunity is initiated. The *Cf* resistance genes encode leucine-rich repeat (LRR)-RLPs conferring immunity to specific races of the pathogenic fungus *Cladosporium fulvum* causing leaf mold disease of tomato (*Solanum lycopersicum*) (Joosten & de Wit, 1999; Rivas & Thomas, 2005; Stergiopoulos & de Wit, 2009). Distinct members of the Cf receptor family are activated upon recognition of their matching ligands, which are the so-called avirulence (Avr) proteins that are secreted by the fungus upon invasion of tomato leaves. The founding member of the Cf proteins and LRR-RLPs in general is Cf-9, which mediates resistance to Avr9-producing strains of *C. fulvum*, whereas for example Cf-2 and Cf-4 recognize Avr2 and Avr4, respectively (Rivas & Thomas, 2005). The Cf receptors mediate race-specific immunity to *C. fulvum*, which is typically associated with the hypersensitive response (HR), a form of programmed cell death.

Well-characterized PRRs are FLAGELLIN SENSING 2 (FLS2) and ELONGATION FACTOR-TU RECEPTOR (EFR), which are LRR-RLKs initiating broad-spectrum immunity against bacterial infection upon detection of the pathogen-associated molecular patterns (PAMPs) flagellin (flg22) and EF-Tu, respectively (Boller & Felix, 2009). To activate immune signaling, FLS2 and EFR form ligand-induced complexes with members of the SOMATIC EMBRYOGENESIS RECEPTOR KINASE (SERK) LRR-RLK family, of which BRI1-ASSOCIATED KINASE 1 (BAK1)/SERK3 plays a prominent role in FLS2-mediated immunity (Heese *et al*., 2007; Roux *et al*., 2011). BAK1 was initially found to interact with the LRR-RLK brassinosteroid receptor BRASSINOSTEROID INSENSITIVE 1 (BRI1), and is involved in a number of immune pathways and developmental processes, in line with its broader function as a co-receptor (Santiago *et al*., 2013; Sun *et al*., 2013; Liebrand *et al*., 2014). All receptors mentioned above are plasma membrane-localized proteins allowing them to detect extracellular microbe-derived ligands. In order to reach their destination after their maturation in the endoplasmic reticulum (ER), they have to be targeted to the plasma membrane by secretory trafficking (Beck *et al*., 2012a). As part of the immune response, plasma membrane-resident FLS2, when activated by flg22, is recruited to ARA7/RabF2b- and ARA6/RabF1-positive endosomes, which are compartments of the late endosomal trafficking pathway (Robatzek *et al*., 2006; Beck *et al*., 2012c). This process depends on BAK1/SERK3 (Chinchilla *et al*., 2007; Beck *et al*., 2012c), which itself undergoes constitutive endocytosis (Russinova *et al*., 2004).

In contrast to RLKs, RLPs lack an intracellular signaling domain and therefore it has been long suggested that these receptors require the interaction with a protein kinase to initiate signaling (Jones *et al*., 1994; Joosten & de Wit, 1999; Liebrand *et al*., 2014). For example, the Toll-like receptors (TLRs) of the mammalian innate immune system require interaction with the MyD88 adaptor protein to recruit the kinase IRAK and subsequently activate immune signaling (Moresco *et al*., 2011; Broz & Monack, 2013). Following the paradigm of the involvement of BAK1/SERK3 in RLK-mediated signaling, there is genetic evidence for a role of BAK1/SERK3 also in plant defense responses mediated by RLPs. Both resistance to the fungal pathogen *Verticillium dahliae*, conferred by the tomato LRR-RLP Ve1 and immunity elicited by *Sclerotinia sclerotiorum* through *Arabidopsis thaliana* RLP30, are dependent on BAK1/SERK3 (Fradin *et al*., 2009; Fradin *et al*., 2011; Zhang *et al*., 2013). Furthermore, another SERK family member, SERK1, was also shown to be genetically required for Ve1 and Cf-4 antifungal immunity in *Arabidopsis* and tomato, respectively (Fradin *et al*., 2011). However, no molecular interaction between these LRR-RLPs and the SERKs has been reported, leaving the possibility that the observed phenotypes are indirect, for example through BAK1/SERK3 function in cell death control or damage-induced immunity (Kemmerling *et al*., 2007; Gao *et al*., 2009; Schwessinger *et al*., 2011; Liu *et al*., 2013; Tintor *et al*., 2013). In addition to its positive regulatory role, BAK1/SERK3 was also described to negatively regulate immunity in tomato (*Sl*) mediated by *Sl*Eix2, an LRR-RLP recognizing fungal xylanase (Bar *et al*., 2010). In a BAK1/SERK3-dependent manner, the close homologue *Sl*Eix1 impairs *Sl*Eix2 endocytosis and HR (Bar *et al*., 2010).

Besides genetic evidence for a role of SERK1, no direct involvement in Cf-mediated immunity of the SERKs has been reported to date. Instead, another LRR-RLK, referred to as SUPPRESSOR OF BIR1-1/EVERSHED (SOBIR1/EVR), has recently been identified as a critical component in LRR-RLP-mediated immunity (Jehle *et al*., 2013; Liebrand *et al*., 2013; Zhang *et al*., 2013; Zhang *et al*., 2014). *Sl*SOBIR1 interacts specifically with RLPs, including the Cf-2, Cf-4 and Cf-9 proteins, and this interaction occurred independently of the presence of their matching Avr ligands. The constitutive association of these LRR-RLPs with *Sl*SOBIR1 is required for RLP stability and has been proposed to allow activation of a cytoplasmic signaling cascade upon perception of an Avr by the interacting Cf protein (Liebrand *et al*., 2013; Liebrand *et al*., 2014). It however remains to be elucidated how ligand-dependent activation of LRR-RLP signaling actually occurs.

Colonization of tomato leaf tissue by *C. fulvum* is fully confined to the apoplast and the fungus does not form specialized feeding structures like haustoria. Consistent with this life-style, the Avr proteins secreted by the fungus appear to solely accumulate in the apoplast (Joosten & de Wit, 1999; Thomma *et al*., 2005; Stergiopoulos & de Wit, 2009; Joosten, 2012). This suggests that the Cf receptors perceive the various secreted Avr proteins of *C. fulvum* at the plasma membrane of the host cells. However, the exact subcellular localization of Cf proteins has remained unclear ever since their first identification (Jones *et al*., 1994). Initially it was shown that Cf-9 is present both at the ER and at the plasma membrane (Piedras *et al*., 2000). A subsequent study demonstrated a role of a C-terminal dilysine motif of Cf-9 in ER targeting (Benghezal *et al*., 2000), which however was later found to be dispensable for Cf-9 function (van der Hoorn *et al*., 2001). More recently, ER-resident chaperones were identified as interacting proteins of Cf-4 and were shown to be important for Avr4-triggered immunity (Liebrand *et al*., 2012), which provided further evidence for ER localization of these RLPs.

Here, we have used live-cell imaging of transiently and stably expressed fluorescent protein fusions in *N. benthamiana* to investigate the subcellular localization and dynamics of Cf-4 *in planta*. Our data suggest that Cf-4, together with *Sl*SOBIR1, perceives Avr4 at the plasma membrane of the host cells, a process that subsequently induces the association of Cf-4 with BAK1/SERK3, and likewise with SERK1. We furthermore demonstrate a role of these SERK family members in Cf-mediated immunity and show that the Cf-4 receptor undergoes BAK1/SERK3-dependent endocytosis upon Avr4 perception.

## Materials and Methods

### Plant materials and constructs

*Nicotiana benthamiana* plants were grown under 16 hours of light at 24°C and 45 - 65 % humidity. Various constructs used in this study have been described before (in all cases GFP refers to enhanced (e)GFP); Cf-4-GFP, Cf-9-GFP, *Sl*SOBIR1-GFP, *Sl*SOBIR1-like-GFP, *At*SOBIR1-GFP, *At*SOBIR1-Myc and *At*SOBIR1^D489N^-Myc (Liebrand *et al*., 2012; Liebrand *et al*., 2013); FLS2-YFPn/c (Frei dit Frey *et al*., 2012); MEMB12-mCherry (Geldner *et al*., 2009); VHA-a1-RFP, RFP-ARA7 and ARA6-RFP (Dettmer *et al*., 2006); AtBAK1 (Schwessinger *et al*., 2011). Cf-4-YFPn/c and *Sl*SOBIR1-YFPn/c were obtained by PCR-amplifying the coding regions of the respective genes, which were cloned into the pENTR/D-TOPO^®^ system (Invitrogen) and subsequently recombined into *pAM-PAT-GWPro_35S_* (GenBank AY436765). The *35S*::*ACA8-mCherry* construct was generated by Gateway recombination (Invitrogen) using pDONR201 containing the coding sequences of *ACA8* devoid of stop codon (Frei dit Frey *et al*., 2012) and a *35S::GW-mCherry* destination vector (provided by R. Panstruga, RWTH Aachen University, Germany). *Sl*SOBIR1-HA was obtained by recombining pENTR/D-TOPO^®^-*Sl*SOBIR1 (Liebrand *et al*., 2013) to pGWB14, carrying the HA coding sequence (Nakagawa *et al*., 2007). *Sl*SERK1-Myc and *Sl*SERK3a-Myc were obtained by recombining pENTR/D-TOPO^®^-*Sl*SERK1 and pDONR201^®^-*Sl*SERK3a (Liebrand *et al*., 2013) to pGWB20 carrying the Myc coding sequence (Nakagawa *et al*., 2007), respectively. pUB::*At*BAK1^D416N^ was obtained by PCR amplification from pGWB14-BAK1^D416N^-HA (Schwessinger *et al*., 2011) using primers BAK1-TOPO-F (5’-CAC CAT GGA ACG AAG ATT AAT GAT C-3’) and BAK1-TOPO-R (5’-TTA TCT TGG ACC CGA GGG GTA TTC-3’) introducing a stop codon, directional cloning into pENTR/D-TOPO^®^ and subsequent recombination of pENTR/D-TOPO-BAK1^D416N^ with pUB-DEST (Grefen *et al*., 2010) All recombinations were performed using the classical Gateway LR clonase reaction (Invitrogen). The correctness of all constructs was confirmed by sequencing.

### Transient, stable transformation, and Virus-Induced Gene Silencing (VIGS)

Transient transformation of *N. benthamiana* epidermal cells was done as described before (Choi *et al*., 2013). Briefly, two-day cultures of *Agrobacterium tumefaciens* GV3101 carrying the respective expression constructs in liquid LB medium supplemented with antibiotics were washed in water prior to *N. benthamiana* leaf infiltrations. For single protein localization and co-localization purposes, final OD_600_=0.25 and OD_600_=0.5, respectively, were used for agro-infiltrations. Microscopy was performed at 2-3 dpi. *N. benthamiana* lines stably expressing Cf-4-GFP were obtained by incubating *N. benthamiana* leaf explants with *Agrobacterium* strain GV3101 carrying plasmid pBIN-KS-35S::Cf-4-GFP (Liebrand *et al*., 2012). Selection of transformed plants was done as described before (Horsch *et al*., 1985; Gabriëls *et al*., 2006). Using segregation analysis based on kanamycin resistance, a single locus insertion line was selected. Tobacco Rattle Virus (TRV)-mediated Virus-Induced Gene Silencing (VIGS) in *N. benthamiana* was performed as described before (Liebrand *et al*., 2012). Briefly, *A. tumefaciens* GV3101 carrying *TRV-RNA1* at OD_600_ = 0.4 and GV3101 carrying *TRV-RNA2* containing the respective target sequences at OD_600_ = 0.2 were mixed and co-infiltrated into *N. benthamiana* leaves of two-weeks old plants, and three weeks later leaves were used for further analysis. *TRV* alone was used as a control. The effects on Cf-4-GFP accumulation of VIGS targeting various endogenous *N. benthamiana* genes in combination with co-expression of *A. thaliana* genes were observed via immunoblotting and RT-PCR (Supporting Information Figs. 6c, 9c,d).

### Bioassay for Avr4-induced immunity

TRV constructs targeting *NbSERK3a/b* (Chaparro-Garcia *et al*., 2011), TRV2::*NbSOBIR1/NbSOBIR1-like* (Liebrand *et al*., 2013) and TRV2::*Cf-4* (Gabriëls *et al*., 2006), alongside with TRV2::*GUS* (Tameling & Baulcombe, 2007) controls, were agro-infiltrated into leaves of one-week-old *N. benthamiana:Cf-4-GFP* plants at O.D.600 = 0.5. After three weeks, Avr4 (O.D.600 = 0.03, twice) (Gabriëls *et al*., 2007), RxD460V (O.D.600 = 0.1) (Bendahmane *et al*., 2002) and BAX (O.D.600 = 0.5) (Lacomme & Santa Cruz, 1999), were agro-infiltrated into mature, fully expanded leaves of the TRV-inoculated plants. HR scores were recorded three days after agro-infiltration. In addition, three-week old *NbSERK3a/b*-silenced leaves were transiently transformed with Cf-4-GFP and 3 days later infiltrated with 300μM Avr4 protein. HR was observed 6 days after Avr4 infiltration. VIGS in tomato, followed by inoculation with conidia of *C. fulvum* race 5-pGPD:*GUS* and subsequent GUS staining and quantification were performed as described before (Liebrand *et al*., 2013).

### Immunoblot analysis and co-immunoprecipitation

Immunoblot analysis with anti-GFP and anti-Myc antibodies and pull-down experiments were carried out as previously reported (Liebrand *et al*., 2012; Liebrand *et al*., 2013), with the following modifications: Transiently transformed *N. benthamiana* leaves were infiltrated with Avr4, Avr2, or Avr9 proteins or flg22 peptide, at the indicated concentrations, and total proteins were extracted. For detection of the HA-epitope tag the anti-HA-Biotin High Affinity Antibody (clone 3F10; Roche Applied Science) was used. In contrast to the theoretical mass of Cf-4-GFP (app. 123 kDa) Immunoblot analysis revealed a specific band at about 140 kDa in accordance with what was observed in previous studies (Liebrand *et al*., 2012).

### qRT-PCR analysis

For qRT-PCR, total RNA was isolated from *N. benthamiana:Cf-4-GFP* leaf material, at 2 weeks after agro-inoculation with the various VIGS constructs. RNA extraction, cDNA synthesis and qRT-PCR were performed as described (Liebrand *et al*., 2012). *NbSERK3a/b* expression was investigated using primers rbo16 (5’-TGC GCT GAA GAC CAA CTT GGC T-3’) and rbo17 (5’-CTG AAG CTT GCC CAA TGT GTC G-3’). *NbSERK1* expression was investigated using primers rbo20 (5’-ATT GCA CAG TCT GCG T-3’) and rbo21 (5’-CGA AGG AAT CTC AAT TTA GTC-3’). Expression of *NbSOBIR1* was investigated using primers to266 and to267 (Liebrand *et al*., 2013). Expression of endogenous actin was used to calibrate the expression level of the query genes, as previously described (Liebrand *et al*., 2012). qRT-PCRs to determine the expression levels of *AtBAK1* and *AtBAK1*^*D416N*^ were performed using primers *At*BAK1-F (5’-TGG ACT TGC AAA ACT CAT GG-3’) and *At*BAK1-R (5’-GAT CAAAAG CCC TTT GTC CA-3’).

### Confocal microscopy and image analysis

Confocal laser microscopy was performed using a Leica SP5 laser point scanning microscope (Leica, Germany) mounted with hybrid detectors (HyD™) as described previously (Beck *et al*., 2012c). Briefly, GFP and RFP/mCherry fluorophores were excited using the 488-nm argon laser and the 561 nm diode, respectively, and fluorescence emission was captured between 500 and 550 nm for GFP and between 580 and 620 nm for RFP/mCherry. For GFP only images, chloroplast autofluorescence was captured between 700 and 800 nm. Sequential scan mode was used for simultaneous imaging of GFP and RFP/mCherry. Abaxial sides of *N. benthamiana* leaf discs were imaged. Images were taken using a 63X water immersion objective and processed using the Leica LAS-AF and FIJI (ImageJ) software packages. Spot detection and quantification on confocal micrographs were performed using EndoQuant, which is a modification of EndomembraneQuantifier suitable for standard confocal images (Beck *et al*., 2012c).

## Results

### Cf-4 interacts with SlSOBIR1 at the plasma membrane

To address the localization and dynamics of the Cf-4 receptor in antifungal immunity, we used functional fluorescently-tagged Cf-4 (Cf-4-GFP) mediating Avr4-triggered HR in *N. benthamiana* (Supporting Information Fig. S1; (Liebrand *et al*., 2012)). We monitored Cf-4-GFP subcellular localization and revealed its presence at the plasma membrane of leaf epidermal cells by co-localizing Cf-4-GFP with the plasma membrane autoinhibitory calcium ATPase ACA8, fused to mCherry (Fig. 1, Supporting Information Fig. S2; (Frei dit Frey *et al*., 2012)). The cell surface localization of this Cf protein is in agreement with the described apoplastic localization of its ligand, Avr4, (Joosten *et al*., 1994; van den Burg *et al*., 2006) and the plasma membrane-localization of its constitutive interactor, SOBIR1 (Fig. 2.; (Leslie *et al*., 2010; Liebrand *et al*., 2013)). *Sl*SOBIR1-GFP, its close homolog *Sl*SOBIR1-like fused to GFP and also *At*SOBIR1-GFP were all detected at the plasma membrane, co-localizing with ACA8-mCherry, and internal vesicles (Figs. 1, 2, Supporting Information Figs. S2, S3, S4).

**Fig. 1.**
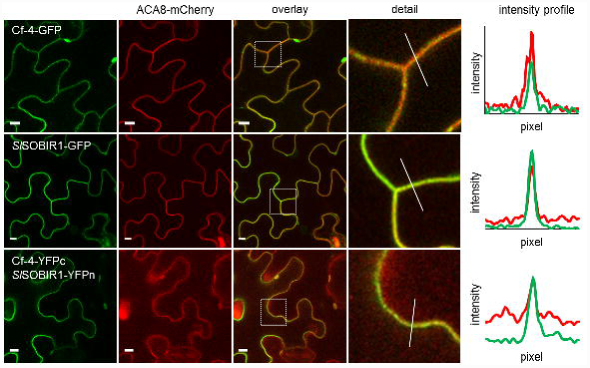
Cf-4 and *Sl*SOBIR1 are present at the plasma membrane. Confocal micrographs show *N. benthamiana* leaf epidermal cells transiently co-expressing the indicated Cf-4 and *Sl*SOBIR1 fusion proteins and plasma membrane-localized ACA8-mCherry. The first panels show GFP/YFP fluorescence, the second panels show mCherry fluorescence, the third panels depict the overlay images of the two fluorescence signals shown in the first and second panels. Overlay images indicate co-localization of the proteins fused to GFP or YFP and mCherry, as a yellow colour is produced. Dashed squares in these images are shown as detail pictures (magnified in the fourth panels). White lines in the detail pictures indicate the ROIs that correspond to intensity profiles in the last panels. Intensity profiles display grey value of pixels across the ROI in the green and red channels, on a scale of 1-300. Images were taken at three days post infiltration (dpi) for Cf-4-GFP and *Sl*SOBIR1-GFP and at two dpi for BiFC of Cf-4, C-terminally fused to the C-terminal half of YFP (YFPc), with *Sl*SOBIR1 C-terminally fused to the N-terminal half of YFP (YFPn); scale bars = 10 μm.

**Fig. 2.**
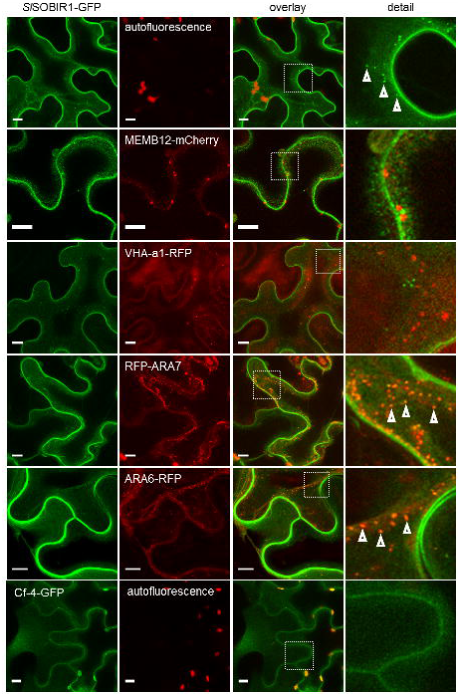
*Sl*SOBIR1 constitutively localizes to endosomes. Confocal micrographs show *N. benthamiana* leaf epidermal cells transiently expressing *Sl*SOBIR1-GFP or Cf-4-GFP (left panels), either without or with co-expression of the indicated organelle markers fused to mCherry/RFP (middle, left panels). Co-localization in the overlay images is depicted by the development of a yellow colour (middle, right panels). Dashed squares in these images are shown as detail pictures (magnified in the right panels). Arrowheads in the detail pictures point at mobile vesicles in the upper right panel and indicate *Sl*SOBIR1 localization at endosomes in the two lower right panels. Images were taken at three dpi; scale bars = 10 μm.

To investigate the subcellular localization of the Cf-4-*Sl*SOBIR1 complex, we used bimolecular fluorescence complementation (BiFC). We observed reconstitution of the YFP molecule by the detection of fluorescence when transiently co-expressing Cf-4 and *Sl*SOBIR1 that were C-terminally fused to respectively the C- and N-terminal halves of YFP (respectively YFPc and YFPn), indicative of Cf-4-*Sl*SOBIR1 heterodimerization (Fig. 1). Cf-4-*Sl*SOBIR1 heterodimerization was found at the plasma membrane, which was confirmed by ACA8-mCherry co-localization (Fig. 1). In this assay, we did not detect positive BiFC when co-expressing FLS2 and *Sl*SOBIR1 fused to the YFP halves, whereas BiFC did occur when co-expressing FLS2-YFPc and FLS2-YFPn (Supporting Information Fig. S5). These observations indicate homodimerization of FLS2 but no heterodimerization between FLS2 and *Sl*SOBIR1, which is consistent with previous findings and thus supporting the specificity of BiFC in this system (Frei dit Frey *et al*., 2012; Sun *et al*., 2012; Liebrand *et al*., 2013). All these results show that Cf-4 localizes at the plasma membrane, where this LRR-RLP interacts with the RLK *Sl*SOBIR1.

### SOBIR1 localizes to endosomes

In addition to its localization at the plasma membrane, and in accordance to what was shown in previous studies, *At*SOBIR1 was also observed at internal vesicles that showed co-labeling with the endocytic tracer FM4-64, indicative of endosomal localization (Leslie *et al*., 2010). To examine the identity of these SOBIR1-positive vesicles, we transiently co-expressed *Sl*SOBIR1-GFP, *Sl*SOBIR1-like-GFP and *At*SOBIR1-GFP with described fluorescent markers of the endomembrane trafficking pathways. These include MEMB12-mCherry for labeling Golgi compartments, VHA-a1-RFP for labeling the *trans*-Golgi network (TGN) and the Rab5 GTPases RFP-ARA7/RabF2b and ARA6/RabF1-RFP for labeling endosomes (Geldner *et al*., 2009; Beck *et al*., 2012c). No co-localization between the SOBIR1-positive vesicles and MEMB12- and VHA-a1-labeled compartments was detected (Fig. 2, Supporting Information Fig. S3a,b). By contrast, we found clear localization of the SOBIR1 receptors to RFP-ARA7/RabF2b- and ARA6/RabF1-RFP-positive endosomes (Fig. 2, Supporting Information Fig. S3c,d). These observations are in agreement with the described FM4-64-positive endosomal localization of *At*SOBIR1-YFP in *Arabidopsis* (Leslie *et al*., 2010), and suggest that constitutive endocytosis of SOBIR1 receptors takes place. Furthermore, co-localization with ARA6/RabF1 indicates that the SOBIR1 receptors might enter the late endosomal pathway (Beck *et al*., 2012c).

### Avr4 triggers endocytosis of the Cf-4/SOBIR1 complex

The observation that Cf-4 interacts with *Sl*SOBIR1 at the plasma membrane, together with the finding that SOBIR1 receptors are present at endosomes, prompted us to investigate whether Cf-4 is endocytosed when activated by Avr4. To test this, we treated *N. benthamiana* leaves transiently and stably expressing Cf-4-GFP with purified Avr4 and Avr2 proteins, of which the latter is specifically recognized by the Cf-2 receptor. When elicited with Avr4, this triggered internalization of Cf-4-GFP (Fig. 3a, Supporting Information Fig. S6). Co-expression with ARA6/RabF1-RFP showed a strong overlap of this marker with the Avr4-induced Cf-4-GFP-positive vesicles, thereby revealing endosomal localization of activated Cf-4 (Fig. 3, Supporting Information Videos S1, S2). Cf-4-As expected, GFP maintained plasma membrane localization upon control treatment with Avr2, indicating that endocytosis of Cf-4 is ligand-specific and depends on the activation of the Cf-4 receptor (Fig. 3a, Supporting Information Fig. S6). Consistent with SOBIR1 localizing to endosomes (Fig. 2, Supporting Information Fig. S3), we observed co-localization of Avr4-induced Cf-4-GFP-positive endosomes and endosomal *Sl*SOBIR1-mCherry (Fig. 3).

**Fig. 3.**
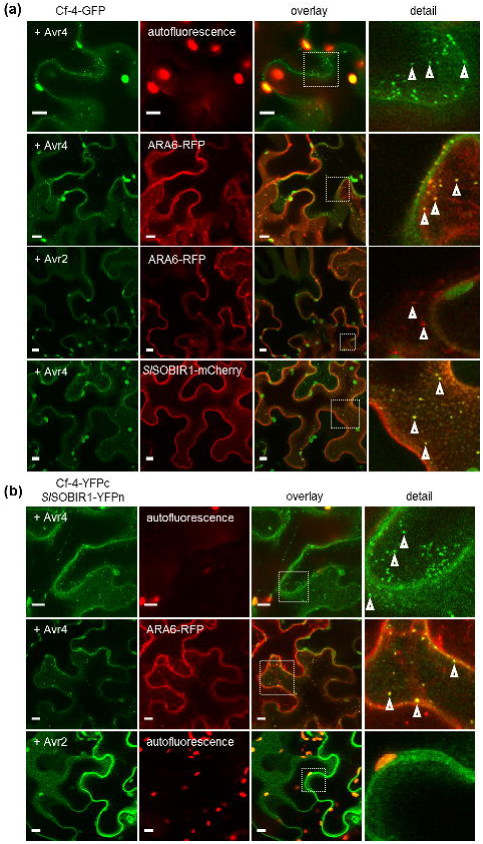
Cf-4 localizes to endosomes in a ligand-dependent manner and together with *Sl*SOBIR1. (**a**) Confocal micrographs show *N. benthamiana* leaf epidermal cells transiently expressing Cf-4-GFP without and with co-expression of ARA6-RFP and *Sl*SOBIR1-mCherry, treated with Avr4 or Avr2 (both at 100μM), as indicated. The left panels show GFP fluorescence, middle left panels show autofluorescence and RFP/mCherry fluorescence, middle right panels depict the overlay images of the two fluorescence signals shown in the left and middle left panels, and the right panels show detail pictures of the dashed squares. Arrowheads point at mobile vesicles positive for Cf-4-GFP and ARA6-RFP and indicate co-localization of Cf-4-GFP with ARA6-RFP and *Sl*SOBIR1-mCherry at endosomes. Images were taken at three dpi and 90 min after elicitation; scale bars = 10 μm. (**b**) Confocal micrographs show *N. benthamiana* leaf epidermal cells transiently expressing Cf-4, C-terminally fused to the C-terminal half of YFP (YFPc) and *Sl*SOBIR1, C-terminally fused to the N-terminal half of YFP (YFPn) (left panels), without or with co-expression of ARA6-RFP (middle left panels) and treated with Avr4 or Avr2 (both at 100μM), as indicated. Co-localization between reconstituted YFP and RFP in the overlay images is depicted by the development of a yellow colour (middle, right panels). Dashed squares in these panels are shown as detail pictures (right panels). Arrowheads point at mobile vesicles in the upper right panel and indicate co-localization at endosomes in the middle right panel. Images were taken at three dpi and 90 min after elicitation; scale bars = 10 μm.

As both, SOBIR1 and Cf-4 underwent endocytosis and localized at late endosomal compartments we questioned whether they might get co-internalized. Therefore we investigated whether Cf-4 is endocytosed together with *Sl*SOBIR1. Transient co-expression of Cf-4 and *Sl*SOBIR1, C-terminally fused to the C- and N-terminal halves of YFP, respectively, results in positive BiFC at the plasma membrane (Fig. 1) and when triggering with Avr4, BiFC was also observed at internal vesicles (Fig. 3b). These vesicles also showed co-localization with ARA6/RabF1-RFP, demonstrating endosomal localization of the reconstituted Cf-4-*Sl*SOBIR1 BiFC heterodimer (Fig. 3b), and in agreement with Avr4-induced co-localization of Cf-4-GFP and *Sl*SOBIR1-mCherry at endosomes (Fig. 3a). Treatment with Avr2 did not trigger re-localization of the plasma membrane-resident reconstituted Cf-4-*Sl*SOBIR1 BiFC heterodimer (Fig. 3b), further supporting ligand-specific internalization of this Cf-4 together with *Sl*SOBIR1. These results indicate that the receptors can be endocytosed as heterodimers, an observation that was previously made for the BRI1-BAK1 complex in *Arabidopsis* protoplasts (Russinova *et al*., 2004). In contrast to this finding, we did not observe flg22-induced endocytosis of the FLS2 homodimer as the reconstituted BiFC signal was maintained at the plasma membrane after elicitation (Supporting Information Fig. S5b). Endocytosis of activated FLS2 depends on BAK1/SERK3 (Chinchilla *et al*., 2007; Beck *et al*., 2012c), and this may indicate that the reconstituted FLS2 BiFC homodimer could be affected in complex formation with BAK1.

### Cf-4 endocytosis requires BAK1/SERK3

Because flg22-activated FLS2 traffics via the ARA6/RabF1-positive late endosomal pathway and depends on functional BAK1/SERK3 (Beck *et al*., 2012c; Choi *et al*., 2013), this raises the possibility that Avr4-induced endocytosis of Cf-4 also involves BAK1/SERK3. To test this, we applied virus-induced gene silencing (VIGS), using a tobacco rattle virus (TRV) construct targeting the two *NbSERK3* homologues (*NbSERK3a* and *b*) in *N. benthamiana* (Chaparro-Garcia *et al*., 2011). Monitoring Cf-4-GFP localization in leaves of *TRV::NbSERK3a/b*-inoculated *N. benthamiana* revealed a strongly reduced amount of Avr4-induced Cf-4-GFP-positive endosomes, as compared to the situation in control leaves from plants that had been inoculated with *TRV* only (Fig. 4, Supporting Information Fig. S10b), while Cf-4 protein levels were unaltered (Supporting Information Fig. S10c). By using a heterologous functional complementation approach, Avr4-induced endocytosis of Cf-4-GFP was partially restored when *AtBAK1* was transiently expressed in *NbSERK3a/b*-silenced leaves. No reduction was observed in the amount of ARA6/RabF1-labelled endosomes when *NbSERK3a/b* was silenced (Fig. 4, Supporting Information Fig. S10b,c). Importantly, this suggests that, as is the case for the RLK FLS2, ligand-induced endocytosis of the RLP Cf-4 is BAK1/SERK3-dependent, and that this RLK plays a role in Cf-4 function. This finding is further strengthened by expression of a kinase-inactive *AtBAK1* variant (*AtBAK1-KD*) in *NbSERK3a/b*-silenced leaves that did not restore Cf-4-GFP endocytosis upon triggering with Avr4 (Fig. 4, Supporting Information Fig. S10b,c).

**Fig. 4.**
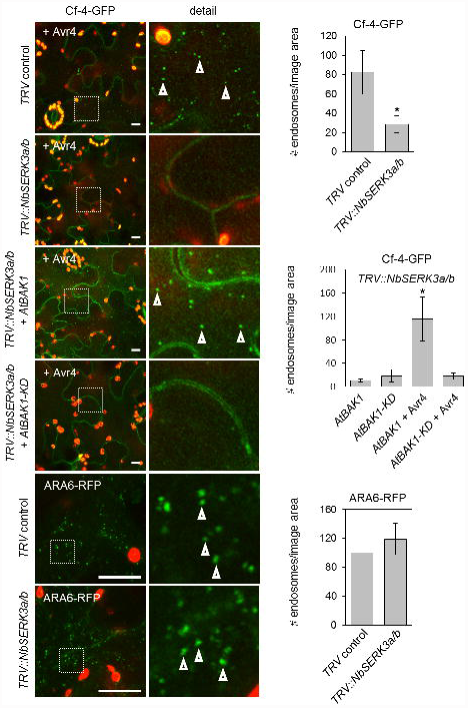
Cf-4 endocytosis requires BAK1/SERK3. Leaves of Cf-4-*N. benthamiana* stable and *N. benthamiana* plants were TRV-silenced for *NbSERK3a/b* and *GUS* as a control for three weeks and subsequently used to transiently express *At*BAK1, *At*BAK1-KD, and ARA6/RabF1-RFP as indicated for three days. Confocal micrographs show Cf-4-GFP localisation upon treatment with Avr4 (100μM, left panels) and ARA6/RabF1-RFP-labelled endosomes (left panels), and detail pictures from dashed squares (middle panels). Arrowheads point at Cf-4-GFP- and ARA6/RabF1-RFP-positive vesicles. Quantification of Cf-4-GFP- and ARA6/RabF1-RFP-positive vesicles was done with EndoQuant (right panels, bars depict means ± 2 SE; n = 6; *p* < 0.05; statistical significant differences are indicated by asterisks). Note, much fewer Cf-4-GFP-positive vesicles are observed in *NbSERK3a/b*-silenced leaves compared to the *GUS*-silenced control. The amount of Cf-4-GFP-positive vesicles increased upon transient co-expression of *At*BAK1 but not when its kinase-inactive variant *At*BAK1-KD was co-expressed. No difference in the amount of ARA6/RabF1-RFP-positive endosomes was observed when *NbSERK3a/b* was silenced. Images were taken 90 min after Avr4 elicitation; scale bars = 10 μm. See Supporting information Fig. S10b,c for transcript abundance of silenced genes and protein levels of Cf-4.

As a next step, we investigated whether SOBIR1 is required for Avr4-induced Cf-4 endocytosis using a similar heterologous functional complementation approach. In agreement with previous findings revealing that Cf-4 abundance is dependent on SOBIR1 (Liebrand *et al*., 2013), Cf-4-GFP levels in *N. benthamiana* were increased by transient expression of *At*SOBIR1-Myc and its kinase-inactive variant (Supporting Information Fig. S7c). Importantly, transient expression of *At*SOBIR1-Myc but not kinase-inactive *At*SOBIR1-KD-Myc restored Avr4-induced endocytosis of Cf-4-GFP in *NbSOBIR1/-like*-silenced leaves (Supporting Information Fig. S7a,b.). Altogether, these data indicate that the kinase activities of both BAK1 and SOBIR1 are required for endocytosis of Cf-4 upon Avr4 recognition, suggesting the active removal of triggered receptors from the plasma membrane.

### Cf-4 associates with SERK members in a ligand-depending manner

We addressed a possible role of BAK1/SERK3 in Cf-4-mediated immunity by co-immunoprecipitation experiments. Cf-4-GFP was purified from *N. benthamiana* leaves transiently co-expressing Cf-4-GFP, *Sl*SOBIR1-HA and *Sl*SERK3a-Myc, that had either been mock-treated or challenged with Avr4, Avr2 or flg22 at two days after agro-infiltration of the three constructs. While C-terminally tagged BAK1/SERK3 fusion proteins are not signalling competent after flg22 and elf18 trigger during immunity their recruitment behavior to the FLS2 receptor complex is still intact, which suggests suitability for interaction studies with other receptors (Ntoukakis *et al*., 2011). In all cases, *Sl*SOBIR1-HA was detected upon immunoprecipitation of Cf-4-GFP, independent of the treatment conditions (Fig. 5, Supporting Information Fig. S8). This confirms the constitutive interaction between these two receptors as shown by BiFC (Fig. 1) and as was previously reported (Liebrand *et al*., 2013). By contrast, *Sl*SERK3a-Myc was only revealed upon immunoprecipitation of Cf-4-GFP from agro-infiltrated leaves that had first been infiltrated with the matching ligand Avr4, but not when infiltrated with Avr2 or flg22, which was included as control treatment (Fig. 5, Supporting Information Fig. S8). Importantly, this demonstrates that ligand-dependent hetero-complex formation of RLPs as the Cf-4 receptor with *Sl*SERK3a takes place, in a way that is similar to what has been shown for RLK-type PRRs (Monaghan & Zipfel, 2012; Liebrand *et al*., 2014).

**Fig. 5.**
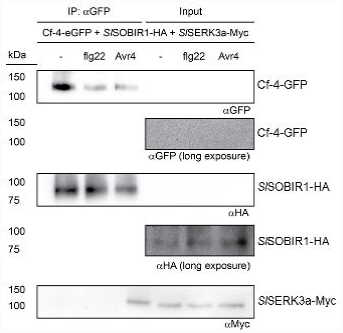
*Sl*SERK3a interacts with Cf-4. Co-immunoprecipitation from GFP-trap bead pull-downs on *N. benthamiana* co-expressing Cf-4-GFP, *Sl*SOBIR1-HA and *Sl*SERK3a-Myc. Leaf samples were taken at two dpi, after 60 min treatments without or with 10μM flg22 or 100μM Avr4, as indicated. Total proteins (Input) and immunoprecipitated proteins (IP) were subjected to SDS/PAGE and blotted. Blots were incubated with αGFP, αHA or αMyc antibodies for the detection of immunoprecipitated Cf-4-GFP and co-purifying *Sl*SOBIR1-HA and *Sl*SERK3a-Myc, respectively.

Because of the previous genetic evidence for a role of *Sl*SERK1 in Cf-mediated immunity (Fradin *et al*., 2011), we examined whether Cf-4 also interacts with this co-receptor by co-immunoprecipitation experiments after transient co-expression of the various receptors in *N. benthamiana* leaves. Unlike what is the case for *Sl*SERK3a-Myc, *Sl*SERK1-Myc was revealed after immunoprecipitation of Cf-4-GFP, independent of elicitation of Cf-4 by Avr4 (Supporting Information Fig. S8). However, the amount of co-immunoprecipitated *Sl*SERK1- Myc was strongly enhanced in the Avr4-treated leaves, as compared to the leaves treated with Avr2. We cannot exclude the possibility that this differential pattern between Cf-4 interaction with *Sl*SERK1 and *Sl*SERK3a is caused by the higher steady state expression levels of *Sl*SERK1-Myc compared to those of *Sl*SERK3a-Myc (Supporting Information Fig. S8; see input). However, following the recent notion of the presence of BRI1-BAK1/SERK3 preassembled heterodimers (Bücherl *et al*., 2013), our data could also indicate that Cf-4 exists in a preformed complex with *Sl*SOBIR1 and *Sl*SERK1, which becomes stabilized and/or recruits increased amounts of *Sl*SERK1 upon elicitation with Avr4.

We next explored the possibility that ligand-induced recruitment of *Sl*SERK1 and -3 could exist as a more general mechanism of Cf receptor complex activation. To this end, we purified Cf-9-GFP from *N. benthamiana* leaves transiently co-expressing this RLP together with *Sl*SOBIR1-HA and either *Sl*SERK1-Myc or *Sl*SERK3a-Myc. Similar to what we observed for activated Cf-4, both *Sl*SERK1-Myc and *Sl*SERK3a-Myc were strongly co-immunoprecipitated with Cf-9-GFP when elicited with its matching ligand effector Avr9 but, as expected, not significantly with Avr4 that was included as control treatment (Supporting Information Fig. S9). This supports the idea that additional SOBIR1-dependent RLPs, such as Ve1, RLP30 and ReMax (Fradin *et al*., 2009; Fradin *et al*., 2011; Zhang *et al*., 2013), also recruit members of the SERK family to form active receptor complexes.

### SERK members mediate Avr4-triggered immunity

Avr4 triggers HR in Cf-4-expressing *N. benthamiana*, which can be diminished by silencing of the gene encoding the Cf-4 receptor itself or *NbSOBIR1/-like* ((Liebrand *et al*., 2013); Fig. 6a). To determine whether BAK1/SERK3 is involved in Cf-4-mediated immunity, we examined the Avr4-triggered HR in Cf-4-GFP-expressing *N. benthamiana* leaves (transiently and stably), without and with silencing of *NbSERK3a/b*. We consistently found that the Avr4-triggered HR was significantly reduced in *NbSERK3a/b*-silenced Cf-4-GFP-expressing leaves, as compared to the leaves of *TRV* control plants (Fig. 6a, Supporting Information Fig. S10a). *NbSERK3a/b*-silencing did not generally affect programmed cell death, as this was still induced by the auto-active variant of the nucleotide binding-LRR immune receptor RxD460V and the pro-apoptotic factor BCL2-ASSOCIATED PROTEIN X (BAX) (Lacomme & Santa Cruz, 1999; Bendahmane *et al*., 2002). The capability of the *NbSERK3a/b*-VIGS construct to knock down the expression of both *NbSERK3a/b* gene homologues was previously shown (Chaparro-Garcia *et al*., 2011). We further investigated the specificity of *NbSERK3a/b*-silencing within the SERK family and found that also a likely *NbSERK1* homologue was knocked down upon expression of the *NbSERK3a/b*-VIGS construct (Supporting Information Fig. S10b). Thus, it could be that Avr4-triggered HR in *N. benthamiana* requires not only BAK1/SERK3 but also SERK1, an observation that is consistent with the earlier finding that VIGS of *SlSERK1* compromises Cf-4-mediated immunity to *C. fulvum* in tomato (Fradin *et al*., 2011).

**Fig. 6.**
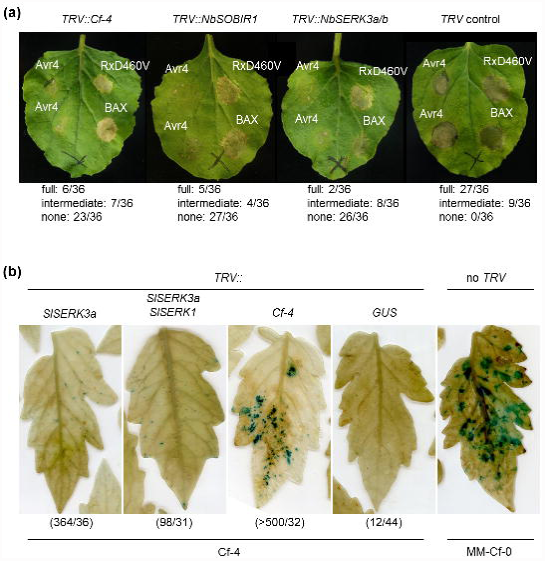
*Sl*SERK3a is required for Avr4-triggered HR and immunity against *C. fulvum*. (**a**) Images show leaves of transgenic *N. benthamiana* plants stably expressing Cf-4-GFP that had been *TRV*-silenced for either *Cf-4*, *NbSOBIR1*, *NbSERK3a/b* or for *GUS* as control, and were subsequently agro-infiltrated with Avr4 (O.D.600 = 0.03), RxD460V (O.D.600 = 0.1) and BAX (O.D.600 = 0.5) as indicated. Images were taken three days after agro-infiltration. HR is observed as brownish cell death. The numbers below the panels indicate the occurrence of full HR, intermediate or no symptoms out of 36 Avr4 agro-infiltrations that were performed. (**b**) Images show leaves two weeks after inoculation with an Avr4-secreting, *GUS*-transgenic strain of *C. fulvum* of MM-Cf-0 tomato as a control, and Cf-4 tomato that had been inoculated with recombinant *TRV* constructs targeting *Cf-4*, *SlSERK3, SlSERK1* or *GUS* three weeks earlier. To visualize *C. fulvum* colonization, leaves were stained for GUS activity. The amount of successful colonization attempts (blue spots) versus the total amount of Cf-4 leaves that were sampled is indicated between parentheses.

To examine whether BAK1/SERK3, in addition to SERK1, is also involved in Cf-4-mediated resistance to Avr4-secreting *C. fulvum* strains, their encoding genes were silenced in Cf-4-expressing tomato plants. A strain secreting Avr4 and expressing the *GUS* reporter gene exhibited strong colonization of tomato leaves lacking the Cf-4 receptor (MM-Cf-0) and when the *Cf-4* gene was silenced (Fig. 6b). Colonization by this fungus was significantly more abundant in Cf-4-expressing tomato leaves silenced for *SlSERK3*, as increased numbers of successful colonization attempts were observed compared to the *GUS*-silenced negative control. This phenotype is reminiscent of the effect of silencing of *SlSOBIR1(-like)* on Cf-4-mediated resistance to *C. fulvum* (Liebrand *et al*., 2013). We furthermore silenced *SlSERK3* together with *SlSERK1*, which also compromised resistance to the pathogen. Taken together, these results show that BAK1/SERK3 is a key positive regulator of full Cf-4-mediated immunity induced upon Avr4 perception.

## Discussion

The Cf signaling pathway is essential for the immune response of tomato to *C. fulvum* (Rivas & Thomas, 2005; Stergiopoulos & de Wit, 2009). A number of Cf signaling components were identified through genetic and proteomic approaches, but the mechanism by which Cf-4 initiates downstream signaling remained unclear (Liebrand *et al*., 2014). Here, we show that both SERK1 and BAK1/SERK3 are recruited to the Cf-4 receptor in an Avr4-dependent manner, a mechanism we additionally confirmed for Avr9-induced activation of Cf-9. Consistently, we show that Cf-4 requires these RLKs for its function. Our observations imply that Avr4 induces the formation of a complex of Cf-4 and *Sl*SERK1/3 to induce Cf-4 signaling. Furthermore, we confirm that Cf-4 interacts with SOBIR1 at the plasma membrane and, more importantly demonstrate ligand-induced SERK3 dependent late endocytic trafficking of the Cf-4 RLP together with *Sl*SOBIR1 as a novel pathway in Cf-mediated downstream events.

The requirement of BAK1/SERK3 hints at clear similarities between the Cf-4/9 effector receptors and FLS2 MAMP receptor pathways, further evidenced by overlapping transcriptional reprogramming upon activating FLS2- and Cf-mediated immune responses (Navarro *et al*., 2004). This overlap potentially involves the regulation of downstream responses through similar components which, in addition to the SERK family members, include E3 ligases (PUB12/13 for FLS2 and CMPG1 for Cf-4 immunity; (Gilroy *et al*., 2011; Lu *et al*., 2011)) and receptor-like cytoplasmic kinases (BIK1/PBL1 for FLS2, ACIK1 for Cf-4 and CAST AWAY for SOBIR1 (EVR) signaling; (Burr *et al*., 2011; Monaghan & Zipfel, 2012; Liebrand *et al*., 2014)). Given these similarities, it is conceivable that the constitutive association between the RLP Cf-4 and the RLK *Sl*SOBIR1 represents a PRR complex, in which the Cf-4 ectodomain mediates specific ligand recognition and the *Sl*SOBIR1 kinase domain is regarded as the signaling part. This is in contrast to PRRs exemplified by FLS2, in which both functions are present within the same molecule. Ligand-induced interaction of PRRs with SERK member RLKs is subsequently required to trigger downstream signaling by both RLP- and RLK-type PRRs (Fig. 7).

**Fig. 7.**
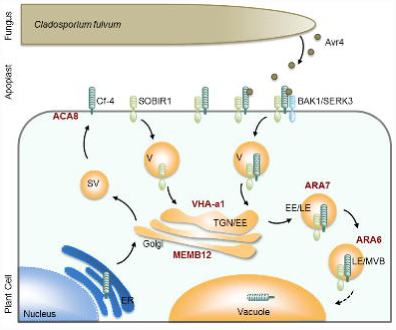
Model of the Cf-4, SOBIR1 and SERK3 subcellular trafficking pathways. Following its folding and maturation in the endoplasmic reticulum (ER), Cf-4 is predominantly trafficked to the plasma membrane, where it constitutively interacts with SOBIR1, which itself is constitutively endocytosed. Upon colonization of Cf-4 tomato leaves, *C. fulvum* secretes Avr4 into the apoplast, which is recognized by Cf-4 and induces interaction of Cf-4 with BAK1/SERK3. This complex is required for Cf-4-mediated immunity (not shown) and endocytosis into ARA7- and ARA6-positive compartments of the late endosomal pathway, destined for vacuolar degradation. Subcellular localization of markers (ACA8, MEMB12, VHA-a1, ARA6 and ARA7) used in this study is depicted in brown lettering. SV, secretory vesicle; V, vesicle; TGN, *trans*-Golgi network; EE, early endosome; LE, late endosome; MVB, multivesicular body.

Regarding Cf-4 as a PRR is in agreement with the observation that Cf-4 not only recognizes the Avr4 protein secreted by *C. fulvum*, but is also activated by the Avr4 homologue produced by the banana pathogen *Mycosphaerella fijiensis* (Stergiopoulos *et al*., 2010). Both *C. fulvum* and *M. fijiensis* Avr4 proteins bind to chitin (van den Burg *et al*., 2003; Stergiopoulos *et al*., 2010) and it was suggested that, reminiscent of PAMP recognition, the chitin-binding motif of Avr4 is the pattern that is recognized through interaction with the LRRs of Cf-4 (Thomma *et al*., 2011). As elicitation with Avr4 appears to maintain the Cf-4 association with *Sl*SOBIR1 ((Liebrand *et al*., 2013); Fig. 5) and induces the interaction of Cf-4 with *Sl*SERK1/3, this could suggest the possibility that a complex consisting of Cf-4, *Sl*SOBIR1 and *Sl*SERK3a is formed. In turn, this could result in increased phosphorylation events between the *Sl*SOBIR1 and *Sl*SERK1/3 kinases, which then trigger downstream signaling (Fig. 7), similar to what has been proposed for the FLS2-BAK1 model (Schwessinger *et al*., 2011). Furthermore, our data show that *Sl*SERK1 and BAK1/*Sl*SERK3 are also recruited to the Cf-9 receptor upon its activation with Avr9, supporting the idea of a general mechanism by which SOBIR1-dependent RLPs form active receptor complexes. Although no molecular interactions have been reported so far, a similar scenario could be proposed for RLP30, which also genetically depends on SOBIR1 and BAK1/SERK3 (Zhang *et al*., 2013). A role for SERK receptors is less clear for *Sl*Eix2, because in this case BAK1 was found to negatively regulate this receptor through interaction with its close homologue *Sl*Eix1 (Bar *et al*., 2010). Future studies should determine whether, similar to the trans-phosphorylation of FLS2 by BAK1/SERK3 (Schwessinger *et al*., 2011), the phosphorylation status of the kinase domain of SOBIR1 also alters upon SERK1/3 recruitment.

Receptor-mediated endocytosis is part of the eukaryotic immune response and for example is found for the FLS2 receptor (Husebye *et al*., 2006; Robatzek *et al*., 2006; Spallek *et al*., 2013). An important role of receptor-mediated endocytosis is to control the abundance of receptor (complexes) at the plasma membrane, a process that is well established in animals and involves lysosomal/vacuolar degradation (Lemmon & Schlessinger, 2010). Activated FLS2 traffics into the late endosomal pathway and is a cargo of multivesicular bodies localizing to the lumen of these late endosomes for delivery to the vacuole (Beck *et al*., 2012c; Spallek *et al*., 2013). This pathway could be responsible for the flg22-induced FLS2 degradation, because chemicals affecting endosomal trafficking inhibit the degradation of this receptor (Smith *et al*., 2014). Our co-localization data strongly suggest that activated Cf-4 is also targeted for vacuolar degradation through the late endosomal pathway, consistent with the observation that Cf-4-GFP protein levels are reduced upon Avr4 elicitation (Supporting Information Fig. S1a). Endosomal sorting for vacuolar degradation requires the transfer of ubiquitin to the plasma membrane cargo and subsequent deubiquitination of the cargo at multivesicular bodies (Beck *et al*., 2012c). The ubiquitin E3 ligase CMPG1 and the deubiquitinating enzyme UBP12 are known positive and negative regulators of Cf-4- and Cf-9-mediated HR, respectively (Ewan *et al*., 2011; Gilroy *et al*., 2011), and given their biochemical function could also be involved in the endocytosis and degradation of Cf proteins.

Ligand-induced activation triggers the formation of a hetero-complex consisting of Cf-4 and *Sl*SERK1/3. This differs from the situation with TLRs from mammals, which recruit cytoplasmic kinases to initiate signaling, but in plants is reminiscent of the FLS2 pathway that also depends on interaction with the co-receptor BAK1/SERK3 to signal. Endocytic removal of activated PRRs, which is also known for TLR4 (Husebye *et al*., 2006), might provide a mechanism to regulate receptor presence at the plasma membrane and associated events (Felix *et al*., 1998; Beck *et al*., 2012a; Smith *et al*., 2014). Understanding how plant cells regulate PRR subcellular localization is essential as pathogens are likely to target components of the trafficking pathway to suppress plant defenses, as was recently found for the *Phytophthora infestans* Avr3a effector (Chaparro-Garcia *et al*.) and bacterial HopM1 (Nomura *et al*., 2011). This latter effector of *Pseudomonas syringae* targets a host ADP ribosylation factor guanine nucleotide exchange factor, referred to as *At*MIN7/BEN1, resulting in its breakdown. *At*MIN7/BEN1 is required for PAMP-triggered immunity and regulates endosomal trafficking possibly at the TGN, where HopM1 can also be found (Nomura *et al*., 2006; Nomura *et al*., 2011). It remains to be seen whether *C. fulvum* also secretes effectors that enter inside plant cells and are able to alter host subcellular trafficking to promote infection success.

## Acknowledgements

We thank the members of the Robatzek laboratory for fruitful discussions, Brande Wulff for reading the manuscript and Cyril Zipfel for providing *At*BAK1 constructs. Zeinu Mussa is acknowledged for help with *N. benthamiana* transformations.

## Supporting Information

**Fig. S1 GFP-tagged Cf-4 migrates with the predicted molecular weight and is functional.** (**a**) Western blot analysis of Cf-4-GFP transiently and stably expressed in *N. benthamiana*, without and with Avr4 (100μM) treatment, as indicated. For transient expression, leaf samples were harvested at three days post infiltration (dpi) and Avr4 elicitation was done for 90 min. Blots were incubated with anti-GFP antibodies for the detection of Cf-4-GFP. CBB, Coomassie Brilliant Blue staining. (**b**) Images of *N. benthamiana* wild type (WT) leaves (top panel) and leaves transiently expressing Cf-4-GFP (bottom panel), treated with either Avr2 or Avr4 (300μM), as indicated. Images were taken at five dpi and at four days after treatment with the Avrs.

**Fig. S2 Subcellular localization of Cf-4, *Sl*SOBIR1-like and *At*SOBIR1.** Confocal micrographs show *N. benthamiana* leaf epidermal cells stably expressing Cf-4-GFP and transiently expressing *Sl*SOBIR1-like-GFP and *At*SOBIR1-GFP as indicated (left panels), and co-expressing plasma membrane-localized ACA8-mCherry (middle panels). Overlay images indicate co-localization of the proteins fused to GFP and mCherry, as a yellow colour is produced (right panels). Images were taken at three dpi; scale bars = 10 μm.

**Fig. S3 *Sl*SOBIR1-like-GFP and *At*SOBIR1-GFP co-localize with endosomal markers.** Confocal micrographs show *N. benthamiana* leaf epidermal cells transiently expressing *Sl*SOBIR1-like-GFP or *At*SOBIR1-GFP (left panels), and co-expressed mCherry/RFP-tagged organelle markers (middle, left panels). Overlay images indicate co-localization through generation of a yellow colour (middle, right panels). Dashed boxes indicated in the middle right panels are depicted as detail pictures on the right (right panels). Arrowheads point at co-localizing endosomes. Images were taken at three dpi; scale bars = 10 μm. (**a**) Co-expression of *Sl*SOBIR1-like-GFP/*At*SOBIR1-GFP with Golgi marker MEMB12-mCherry. (**b**) Co-expression of *Sl*SOBIR1-like-GFP/*At*SOBIR1-GFP with *trans*-Golgi network marker VHA-a1-RFP. (**c**) Co-expression of *Sl*SOBIR1-like-GFP/*At*SOBIR1-GFP with endosome marker RFP-ARA7/RabF2b. (**d**) Co-expression of *Sl*SOBIR1-like-GFP/*At*SOBIR1-GFP with late endosome marker ARA6/RabF1-RFP.

**Fig. S4. Line intensity profiles of Cf-4, SOBIR1 and membrane markers.** Confocal micrographs show transient co-expression of Cf-4-GFP, *Sl*SOBIR1-GFP or Cf-4-YFPc/*Sl*SOBIR1-YFPn with indicated membrane markers and effector treatments. Left panels correspond to confocal micrographs displayed in (**a**) Fig. 1, (**b**) Fig. 2a, (**c**) Fig. 2b. Dashed squares in these panels are shown as detail pictures (magnified in middle panels). White lines in detail pictures indicate the ROIs that correspond to intensity profiles in the last panels. Intensity profiles display grey value of pixels across the ROI in the green and red channels on a scale of 1-300.

**Fig. S5 *Sl*SOBIR1 does not interact with FLS2 in BiFC experiments.** (**a**) Confocal micrographs show *N. benthamiana* leaf epidermal cells transiently co-expressing FLS2 and *Sl*SOBIR1 C-terminally fused to the C- or N-terminal halves of YFP (YFPc and YFPn, respectively), as indicated. Left panels show absence or presence of YFP fluorescence, the middle panels show autofluorescence, and the right panels show the overlay of the fluorescence signals. Images were taken at two dpi, scale bars = 100 μm. (**b**) Confocal micrographs of *N. benthamiana* leaf epidermal cells transiently co-expressing FLS2-YFPc and FLS2-YFPn, either without or with flg22 treatment (10μM) for 90 min, as indicated. Left panels show absence or presence of YFP fluorescence, the middle panels show autofluorescence, and the right panels show the overlay of the fluorescence signals. Images were taken at two dpi, scale bars = 10 μm.

**Fig. S6 Cf-4-GFP endocytosis in stable transgenic plants.** Confocal micrographs of Cf-4-GFP-transgenic *N. benthamiana* plants show leaf epidermal cells expressing Cf-4-GFP, either treated with Avr4 or Avr2 (both at 100 μM), as indicated. Left panels show GFP fluorescence, middle left panels show autofluorescence, and middle right panels show the overlay of the fluorescence signals shown in the left and middle left panels. Arrowheads in the detail pictures (right panels) point at mobile vesicles (top right panel). Images were taken at 90 minutes after elicitation; scale bars = 10 μm.

**Fig. S7 Cf-4 endocytosis requires SOBIR1 kinase activity.** Leaves of Cf-4-GFP transgenic *N. benthamiana* plants that had been TRV-silenced for *NbSOBIR1/-like* were used for transient expression of *AtSOBIR1-Myc* and its kinase-inactive variant *AtSOBIR1-KD-Myc*. (**a**) Confocal micrographs show Cf-4-GFP localisation upon treatment with Avr4 (100μM, left panels) and detail pictures from dashed squares (middle panels). Arrowheads point at Cf-4-GFP-positive vesicles. Images were taken at three weeks after inoculation with *TRV*, at three dpi for transient expression, and at 90 min after elicitation; scale bars = 10 μm. (**b**) Quantification of Cf-4-GFP-positive vesicles was done with EndoQuant (bars depict means ± 2 SE; n = 6; *p* < 0.05; statistical significant differences are indicated by asterisks). Transient co-expression of *At*BAK1, but not its kinase-inactive variant *At*BAK1-KD, increased Cf-4-GFP-positive vesicles in *NbSOBIR1/-like-*silenced leaves. (**c**) Immunoblots from extracted total proteins of Cf-4-GFP-transgenic *N. benthamiana* plants that were TRV-silenced for *NbSOBIR1/-like* and subsequently transiently transformed with *AtSOBIR1-Myc* or *AtSOBIR1-KD-Myc*. Cf-4-GFP, *At*SOBIR1-Myc and *At*SOBIR1-KD-Myc were revealed with αGFP and αMyc antibodies as indicated. Ponceau staining is shown for equal loading.

**Fig. S8 Both *Sl*SERK1 and *Sl*SERK3a interact with the Cf-4.** Co-immunoprecipitation from GFP-trap bead pull-downs on *N. benthamiana* co-expressing Cf-4-GFP and *Sl*SOBIR1-HA, and either *Sl*SERK3a-Myc or *Sl*SERK1-Myc. Leaf samples were taken at two dpi, after 60 min treatment with Avr2 or Avr4 (100μM), as indicated. Total proteins (Input) and immunoprecipitated proteins (IP) were subjected to SDS/PAGE and blotted. Blots were incubated with αGFP, αHA or αMyc antibodies for the detection of immunoprecipitated Cf-4-GFP and co-purifying *Sl*SOBIR1-HA and *Sl*SERK3a-Myc or *Sl*SERK1-Myc, respectively.

**Fig. S9 Avr9 induces interaction of Cf-9 with both *Sl*SERK1 and *Sl*SERK3a.** Co-immunoprecipitation from GFP-trap bead pull-downs on *N. benthamiana* co-expressing Cf-9-GFP and *Sl*SOBIR1-HA, and either *Sl*SERK1-Myc or *Sl*SERK3a-Myc. Leaf samples were taken at two dpi, after 60 min treatment with Avr9 or Avr4 (100μM), as indicated. Total proteins (Input) and immunoprecipitated proteins (IP) were subjected to SDS/PAGE and blotted. Blots were incubated with αGFP, αHA or αMyc antibodies for the detection of immunoprecipitated Cf-9-GFP and co-purifying *Sl*SOBIR1-HA and *Sl*SERK1-Myc or *Sl*SERK3a-Myc, respectively.

**Fig. S10 *Sl*SERK3 is required for Avr4-induced HR.** (**a**) Images show *N. benthamiana* leaves from *TRV::NbSERK3a/b*-inoculated plants and *TRV*-inoculated controls, transiently expressing Cf-4-GFP and treated with Avr4 protein (300μM). Cf-4-GFP was agro-infiltrated at 3 weeks post inoculation with the recombinant *TRV* constructs and one day later the Avr4 protein was infiltrated. Images were taken at six days after Avr4 infiltration. HR is observed as brownish cell death. (**b**) qRT-PCR analysis showing *NbSERK1*, *NbSERK3a/b* and *NbSOBIR1* expression in *N. benthamiana* inoculated with the indicated *TRV* VIGS constructs. Expression of query genes was normalized to endogenous *NbActin* expression levels. Data presented have been combined from twelve (*NbSOBIR1* expression) or 24 (*NbSERK1* and *NbSERK3* expression) individual qRT-PCRs, based on three independent biological replicates, for each gene of which the expression was studied. Statistical differences (α=0.05) between groups were calculated by a one-way ANOVA and different groupings are indicated. Error bars represent the standard deviation. (**c**) Immunoblots from total protein extracts of Cf-4-GFP-transgenic *N. benthamiana* plants that were TRV-silenced for *NbSERK3a/b* and subsequently transiently transformed with *AtBAK1* and *AtBAK1-KD*. Cf-4-GFP was revealed with αGFP. Ponceau staining is shown for equal loading. (**d**) RT-PCR on cDNA obtained from total RNA extracts of Cf-4-GFP-transgenic *N. benthamiana* plants that were TRV-silenced for *NbSERK3a/b* and subsequently transiently transformed with *AtBAK1* and *AtBAK1-KD*.

**Video S1. Mobile vesicles in transient Cf-4-GFP expression.** Time series of confocal micrographs of *N. benthamiana* leaf epidermal cells transiently expressing Cf-4-GFP and treated with Avr4 (100 μM). Images were taken at 3 days post infiltration (dpi) and 1.5 hours after Avr4 treatment. Frame counter is displayed in the top left. Acquisition speed is ∼4 seconds per frame. Scale bar = 10 μm.

**Video S2. Mobile vesicles in stable Cf-4-GFP expression.** Time series of confocal micrographs of *N. benthamiana* leaf epidermal cells stably expressing Cf-4-GFP and treated with Avr4 (100 μM). Images were taken 1.5 hours after Avr4 treatment. Frame counter is displayed in the top left. Acquisition speed is ∼4 seconds per frame. Scale bar = 10 μm.

## References

Bar M, Sharfman M, Ron M, Avni A. 2010. BAK1 is required for the attenuation of ethylene-inducing xylanase (Eix)-induced defense responses by the decoy receptor LeEix1. The Plant Journal 63(5): 791–800.

Beck M, Heard W, Mbengue M, Robatzek S. 2012a. The INs and OUTs of pattern recognition receptors at the cell surface. Current Opinion in Plant Biology 15(4): 367–374.

Beck M, Zhou J, Faulkner C, Mac D, Robatzek S. 2012c. Spatio-temporal cellular dynamics of the *Arabidopsis* flagellin receptor reveal activation status-dependent endosomal sorting. The Plant Cell 24(10): 4205–4219.

Bendahmane A, Farnham G, Moffett P, Baulcombe DC. 2002. Constitutive gain-of-function mutants in a nucleotide binding site-leucine rich repeat protein encoded at the *Rx* locus of potato. The Plant Journal 32(2): 195–204.

Benghezal M, Wasteneys GO, Jones DA. 2000. The C-terminal dilysine motif confers endoplasmic reticulum localization to type I membrane proteins in plants. The Plant Cell 12(7): 1179–1201.

Boller T, Felix G. 2009. A renaissance of elicitors: perception of microbe-associated molecular patterns and danger signals by pattern-recognition receptors. Annual review of plant Biology 60: 379–407.

Broz P, Monack DM. 2013. Newly described pattern recognition receptors team up against intracellular pathogens. Nature Reviews Immunology 13(8): 551–565.

Bücherl ca, van Esse gw, Kruis A, Luchtenberg J, Westphal AH, Aker J, van Hoek A, Albrecht C, Borst JW, de Vries sc. 2013. Visualization of BRI1 and BAK1(SERK3) membrane receptor heterooligomers during Brassinosteroid signaling. Plant Physiology 162(4): 1911–1925.

Burr CA, Leslie ME, Orlowski SK, Chen I, Wright CE, Daniels MJ, Liljegren SJ. 2011. Cast away, a membrane-associated receptor-like kinase, inhibits organ abscission in Arabidopsis. Plant Physiology 156(4): 1837–1850.

Chaparro-Garcia A, Schwizer S, Sklenar J, Yoshida K, Bos JIB, Schornack S, Jones AME, Bozkurt TO, Kamoun S. Phytophthora infestans RXLR-WY effector AVR3a associates with a Dynamin-Related Protein involved in endocytosis of a plant pattern recognition receptor. biorXiv doi: 10.1101/012963.

Chaparro-Garcia A, Wilkinson RC, Gimenez-Ibanez S, Findlay K, Coffey MD, Zipfel C, Rathjen JP, Kamoun S, Schornack S. 2011. The receptor-like kinase Serk3/Bak1 is required for basal resistance against the late blight pathogen *Phytophthora infestans In Nicotiana benthamiana*. PLoS ONE 6(1).

Chinchilla D, Zipfel C, Robatzek S, Kemmerling B, Nürnberger T, Jones JDG, Felix G, Boller T. 2007. A flagellin-induced complex of the receptor FLS2 and BAK1 initiates plant defence. Nature 448(7152): 497–500.

Choi SW, Tamaki T, Ebine K, Uemura T, Ueda T, Nakano A. 2013. RABA members act in distinct steps of subcellular trafficking of the FLAGELLIN SENSING2 receptor. The Plant Cell 25(3): 1174–1187.

Dettmer J, Hong-Hermesdorf A, Stierhof YD, Schumacher K. 2006. Vacuolar H+-ATPase activity is required for endocytic and secretory trafficking in Arabidopsis. The Plant Cell 18(3): 715–730.

Ewan R, Pangestuti R, Thornber S, Craig A, Carr C, O’Donnell L, Zhang C, Sadanandom A. 2011. Deubiquitinating enzymes AtUBP12 and AtUBP13 and their tobacco homologue NtUBP12 are negative regulators of plant immunity. New Phytologist 191(1): 92–106.

Faulkner C, Robatzek S. 2012. Plants and pathogens: Putting infection strategies and defence mechanisms on the map. Current Opinion in Plant Biology 15(6): 699–707.

Felix G, Baureithel K, Boller T. 1998. Desensitization of the perception system for chitin fragments in tomato cells. Plant Physiology 117(2): 643–650.

Fradin EF, Abd-El-Haliem A, Masini L, van den Berg GCM, Joosten MHAJ, Thomma BPHJ. 2011. Interfamily transfer of tomato *Ve1* mediates *Verticillium* resistance *In Arabidopsis*. Plant Physiology 156(4): 2255–2265.

Fradin EF, Zhang Z, Ayala JCJ, Castroverde CDM, Nazar RN, Robb J, Liu CM, Thomma BPHJ. 2009. Genetic dissection of *Verticillium* wilt resistance mediated by tomato Ve1. Plant Physiology 150(1): 320–332.

Frei dit Frey nf, Mbengue M, Kwaaitaal M, Nitsch L, Altenbach D, Häweker H, Lozano-Duran R, Njo MF, Beeckman T, Huettel B, et al. 2012. Plasma membrane calcium ATPases are important components of receptor-mediated signaling in plant immune responses and development. Plant Physiology 159(2): 798–809.

Gabriëls SHEJ, Takken FLW, Vossen JH, De Jong cf, Liu Q, Turk SCHJ, Wachowski LK, Peters J, Witsenboer HMA, de Wit pjgm, et al. 2006. cDNA-AFLP combined with functional analysis reveals novel genes involved in the hypersensitive response. Molecular Plant-Microbe Interactions 19(6): 567–576.

Gabriëls SHEJ, Vossen JH, Ekengren SK, van Ooijen G, Abd-El-Haliem am, van den Berg gcm, Rainey DY, Martin GB, Takken FLW, de Wit pjgm, et al. 2007. An NB-LRR protein required for HR signalling mediated by both extra-and intracellular resistance proteins. The Plant Journal 50(1): 14–28.

Gao M, Wang X, Wang D, Xu F, Ding X, Zhang Z, Bi D, Cheng YT, Chen S, Li X, et al. 2009. Regulation of cell death and innate immunity by two receptor-like kinases *In Arabidopsis*. Cell Host & Microbe 6(1): 34–44.

Geldner N, Dénervaud-Tendon V, Hyman DL, Mayer U, Stierhof YD, Chory J. 2009. Rapid, combinatorial analysis of membrane compartments in intact plants with a multicolor marker set. The Plant Journal 59(1): 169–178.

Gilroy EM, Taylor RM, Hein I, Boevink P, Sadanandom A, Birch PRJ. 2011. CMPG1-dependent cell death follows perception of diverse pathogen elicitors at the host plasma membrane and is suppressed by Phytophthora infestans RXLR effector AVR3a. New Phytologist 190(3): 653–666.

Grefen C, Donald N, Hashimoto K, Kudla J, Schumacher K, Blatt MR. 2010. A ubiquitin-10 promoter-based vector set for fluorescent protein tagging facilitates temporal stability and native protein distribution in transient and stable expression studies. Plant J 64(2): 355–365.

Heese A, Hann DR, Gimenez-Ibanez S, Jones AME, He K, Li J, Schroeder JI, Peck SC, Rathjen JP. 2007. The receptor-like kinase SERK3/BAK1 is a central regulator of innate immunity in plants. Proceedings of the National Academy of Sciences 104(29): 12217–12222.

Horsch RB, Fry JE, Hoffmann NL, Eichholtz D, Rogers SG, Fraley RT. 1985. A simple and general method for transferring genes into plants. Science 227(4691): 1229–1230.

Husebye H, Halaas Ø, Stenmark H, Tunheim G, Sandanger Ø, Bogen B, Brech A, Latz E, Espevik T. 2006. Endocytic pathways regulate Toll-like receptor 4 signaling and link innate and adaptive immunity. EMBO Journal 25(4): 683–692.

Jehle AK, Fürst U, Lipschis M, Albert M, Felix G. 2013. Perception of the novel MAMP eMax from different *Xanthomonas* species requires the *Arabidopsis* receptor-like protein ReMAX and the receptor kinase SOBIR. Plant Signaling & Behavior 8(12): e27408.

Jones DA, Thomas CM, Hammond-Kosack KE, Balint-Kurti PJ, Jones JDG. 1994. Isolation of the tomato *Cf-9* gene for resistance to *Cladosporium fulvum* by transposon tagging. Science 266(5186): 789–793.

Joosten MHAJ. 2012. Isolation of apoplastic fluid from leaf tissue by the vacuum infiltration-centrifugation technique. Methods in molecular biology 835: 603–610.

Joosten MHAJ, Cozijnsen TJ, De Wit PJGM. 1994. Host resistance to a fungal tomato pathogen lost by a single base-pair change in an avirulence gene. Nature 367(6461): 384–386.

Joosten MHAJ, de Wit PJGM. 1999. The tomato-*Cladosporium fulvum* interaction: A versatile experimental system to study plant-pathogen interactions. Annual Review of Phytopathology 37: 335–367.

Kemmerling B, Schwedt A, Rodriguez P, Mazzotta S, Frank M, Qamar SA, Mengiste T, Betsuyaku S, Parker JE, Müssig C, et al. 2007. The BRI1-associated kinase 1, BAK1, has a brassinolide-independent role in plant cell-death control. Current Biology 17(13): 1116–1122.

Lacomme C, Santa Cruz S. 1999. Bax-induced cell death in tobacco is similar to the hypersensitive response. Proceedings of the National Academy of Sciences of the United States of America 96(14): 7956–7961.

Lemmon MA, Schlessinger J. 2010. *Cell signaling by receptor tyrosine kinases*. Cell 141(7): 1117–1134.

Leslie ME, Lewis MW, Youn JY, Daniels MJ, Liljegren SJ. 2010. The EVERSHED receptor-like kinase modulates floral organ shedding in Arabidopsis. Development 137(3): 467–476.

Liebrand TWH, Smit P, Abd-El-Haliem A, De Jonge R, Cordewener JHG, America AHP, Sklenar J, Jones AME, Robatzek S, Thomma BPHJ, et al. 2012. Endoplasmic reticulum-quality control chaperones facilitate the biogenesis of Cf receptor-like proteins involved in pathogen resistance of tomato. Plant Physiology 159(4): 1819–1833.

Liebrand TWH, van den Berg gcm, Zhang Z, Smit P, Cordewener JHG, America AHP, Sklenar J, Jones AME, Tameling WIL, Robatzek S, et al. 2013. The receptor-like kinase SOBIR1/EVR interacts with receptor-like proteins in plant immunity against fungal infection. Proceedings of the National Academy of Sciences of the United States of America 110(24): 10010–10015.

Liebrand TWH, van den Burg HA, Joosten MHAJ. 2014. Two for all: receptor-associated kinases SOBIR1 and BAK1. Trends in Plant Science 19(2): 123–132.

Liu Z, Wu Y, Yang F, Zhang Y, Chen S, Xie Q, Tian X, Zhou JM. 2013. BIK1 interacts with PEPRs to mediate ethylene-induced immunity. Proceedings of the National Academy of Sciences of the United States of America 110(15): 6205–6210.

Lu D, Lin W, Gao X, Wu S, Cheng C, Avila J, Heese A, Devarenne TP, He P, Shan L. 2011. Direct ubiquitination of pattern recognition receptor FLS2 attenuates plant innate immunity. Science 332(6036): 1439–1442.

Monaghan J, Zipfel C. 2012. Plant pattern recognition receptor complexes at the plasma membrane. Current Opinion in Plant Biology 15: 349–357.

Moresco EMY, LaVine D, Beutler B. 2011. Toll-like receptors. Current Biology 21(13): R488–R493.

Nakagawa T, Kurose T, Hino T, Tanaka K, Kawamukai M, Niwa Y, Toyooka K, Matsuoka K, Jinbo T, Kimura T. 2007. Development of series of gateway binary vectors, pGWBs, for realizing efficient construction of fusion genes for plant transformation. Journal of Bioscience and Bioengineering 104(1): 34–41.

Navarro L, Zipfel C, Rowland O, Keller I, Robatzek S, Boller T, Jones JDG. 2004. The transcriptional innate immune response to flg22. Interplay and overlap with Avr gene-dependent defense responses and bacterial pathogenesis. Plant Physiology 135(2): 1113–1128.

Nomura K, Debroy S, Lee YH, Pumplin N, Jones J, He SY. 2006. A bacterial virulence protein suppresses host innate immunity to cause plant disease. Science 313(5784): 220–223.

Nomura K, Mecey C, Lee YN, Imboden LA, Chang JH, He SY. 2011. Effector-triggered immunity blocks pathogen degradation of an immunity-associated vesicle traffic regulator in Arabidopsis. Proceedings of the National Academy of Sciences of the United States of America 108(26): 10774–10779.

Ntoukakis V, Schwessinger B, Segonzac C, Zipfel C. 2011. Cautionary Notes on the Use of C-Terminal BAK1 Fusion Proteins for Functional Studies. The Plant Cell 23(11): 3871–3878.

Piedras P, Rivas S, Dröge S, Hillmer S, Jones JDG. 2000. Functional, c-myc-tagged *Cf-9* resistance gene products are plasma-membrane localized and glycosylated. The Plant Journal 21(6): 529–536.

Rivas S, Thomas CM. 2005. Molecular interactions between tomato and the leaf mold pathogen *Cladosporium fulvum*. Annual Review of Phytopathology 43: 395–436.

Robatzek S, Chinchilla D, Boller T. 2006. Ligand-induced endocytosis of the pattern recognition receptor FLS2 *In Arabidopsis*. Genes and Development 20(5): 537–542.

Roux M, Schwessinger B, Albrecht C, Chinchilla D, Jones A, Holton N, Malinovsky FG, Tör M, de Vries S, Zipfel C. 2011. The *Arabidopsis* leucine-rich repeat receptor-like kinases BAK1/SERK3 and BKK1/SERK4 are required for innate immunity to hemibiotrophic and biotrophic pathogens. The Plant Cell 23(6): 2440–2455.

Russinova E, Borst JW, Kwaaitaal M, Caño-Delgado A, Yin Y, Chory J, de Vries Sc. 2004. Heterodimerization and endocytosis of Arabidopsis brassinosteroid receptors BRI1 and AtSERK3 (BAK1). The Plant Cell 16(12): 3216–3229.

Santiago J, Henzler C, Hothorn M. 2013. Molecular mechanism for plant steroid receptor activation by somatic embryogenesis co-receptor kinases. Science 341(6148): 889–892.

Schwessinger B, Roux M, Kadota Y, Ntoukakis V, Sklenar J, Jones A, Zipfel C. 2011. Phosphorylation-dependent differential regulation of plant growth, cell death, and innate immunity by the regulatory receptor-like kinase BAK1. PLoS Genetics 7(4): e1002046.

Smith JM, Salamango DJ, Leslie ME, Collins CA, Heese A. 2014. Sensitivity to Flg22 Is modulated by ligand-induced degradation and de novo synthesis of the endogenous flagellin-receptor FLAGELLIN-SENSING2. Plant Physiology 164(1): 440–454.

Spallek T, Beck M, Ben Khaled S, Salomon S, Bourdais G, Schellmann S, Robatzek S. 2013. ESCRT-I mediates FLS2 endosomal sorting and plant immunity. PLoS Genetics 9(12): e1004035.

Stergiopoulos I, de Wit PJGM. 2009. Fungal effector proteins. Annual Review of Phytopathology 47: 233–263.

Stergiopoulos I, van den Burg Ha, Ökmen B, Beenen HG, van Liere S, Kema GHJ, de Wit pjgm. 2010. Tomato Cf resistance proteins mediate recognition of cognate homologous effectors from fungi pathogenic on dicots and monocots. Proceedings of the National Academy of Sciences of the United States of America 107(16): 7610–7615.

Sun W, Cao Y, Jansen KL, Bittel P, Boller T, Bent AF. 2012. Probing the *Arabidopsis* flagellin receptor: FLS2-FLS2 association and the contributions of specific domains to signaling function. The Plant Cell 24(3): 1096–1113.

Sun Y, Li L, Macho AP, Han Z, Hu Z, Zipfel C, Zhou J-m, Chai J. 2013. Structural basis for flg22-induced activation of the *Arabidopsis* FLS2-BAK1 immune complex. Science 342(6158): 624–628.

Tameling WIL, Baulcombe DC. 2007. Physical association of the NB-LRR resistance protein Rx with a Ran GTPase-activating protein is required for extreme resistance to *Potato virus X*. The Plant Cell 19(5): 1682–1694.

Thomma BPHJ, Nürnberger T, Joosten MHAJ. 2011. Of PAMPs and effectors: the blurred PTI-ETI dichotomy. The Plant Cell 23(1): 4–15.

Thomma BPHJ, Van Esse HP, Crous PW, De Wit pjgm. 2005. *Cladosporium fulvum* (syn. *Passalora fulva*), a highly specialized plant pathogen as a model for functional studies on plant pathogenic *Mycosphaerellaceae*. Molecular Plant Pathology 6(4): 379–393.

Tintor N, Ross A, Kanehara K, Yamada Y, Fan L, Kemmerling B, Nürnberger T, Tsuda K, Saijo Y. 2013. Layered pattern receptor signaling via ethylene and endogenous elicitor peptides during *Arabidopsis* immunity to bacterial infection. Proceedings of the National Academy of Sciences of the United States of America 110(15): 6211–6216.

van den Burg ha, Harrison SJ, Joosten MHAJ, Vervoort J, de Wit pjgm. 2006. *Cladosporium fulvum* Avr4 protects fungal cell walls against hydrolysis by plant chitinases accumulating during infection. Molecular Plant-Microbe Interactions 19(12): 1420–1430.

van den Burg ha, Westerink N, Francoijs KJ, Roth R, Woestenenk E, Boeren S, de Wit PJGM, Joosten MHAJ, Vervoort J. 2003. Natural disulfide bond-disrupted mutants of AVR4 of the tomato pathogen *Cladosporium fulvum* are sensitive to proteolysis, circumvent Cf-4-mediated resistance, but retain their chitin binding ability. Journal of Biological Chemistry 278(30): 27340–27346.

van der Hoorn RAL, van der Ploeg A, de Wit PJGM, Joosten MHAJ. 2001. The C-terminal dilysine motif for targeting to the endoplasmic reticulum is not required for Cf-9 function. Molecular Plant-Microbe Interactions 14(3): 412–415.

Zhang L, Kars I, Essenstam B, Liebrand TWH, Wagemakers L, Elberse J, Tagkalaki P, Tjoitang D, van den Ackerveken G, van Kan JAL. 2014. Fungal endopolygalacturonases are recognized as microbe-associated molecular patterns by the *Arabidopsis* receptor-like protein RESPONSIVENESS TO BOTRYTIS POLYGALACTURONASES1. Plant Physiology 164(1): 352–364.

Zhang W, Fraiture M, Kolb D, Löffelhardt B, Desaki Y, Boutrot F, Tör M, Zipfel C, Gust AA, Brunner F. 2013. The *Arabidopsis thaliana* receptor-like protein RLP30 and receptor-like kinase SOBIR1/EVR mediate innate immunity toward necrotrophic fungi. The Plant Cell 25(10): 4227–4241.

